# Plant protein-based diets can replace a fish meal-based diet for sustainable growth and body composition of zebrafish

**DOI:** 10.1101/2020.05.18.101733

**Authors:** Toluwalase Anthony Aiyelari, Abdul Shakoor Chaudhry

## Abstract

This 3 × 2 factorial study involving three diets at two stocking densities tested the effect of replacing fish meal (FM) with either soybean meal (SBM) or rapeseed meal (RSM) in diets on growth and body composition of zebrafish (*Danio rerio*). Fish were fed three times daily for eight weeks. Morphometric and water quality parameters were also determined. The survival rate of the fish ranged from 95.2 - 97.8%. The water quality remained within the acceptable limits for tropical aquaculture. The stocking density did not show any significant difference (p>0.05) for the length and weight of the fish. The length, weight and condition factors were significantly higher (p<0.05) in the fish fed FM based diet. The fish length and weight related well (R^2^) across the diets but this was more significant for RSM than those fed the other two diets. The weight gain (WG), feed conversion ratio (FCR) and protein intake (PI) were significantly higher (p<0.05) in the fish fed the FM based diet than the other diets. No significant differences observed (p>0.05) in the specific growth rate (SGR), food intake (FI) and protein efficiency ratio (PER) among the fish fed the three diets. The crude protein CP, nitrogen-free extract (NFE) and ash contents of these fish did not differ significantly (p>0.05). However, the ether extract (EE) of the fish fed SBM diet was significantly lower (p<0.05 than the other two diets. It appears that both SBM and RSM as sustainable source to partially FM in the diets of zebrafish and similar fish species.

## Introduction

Fisheries and aquaculture are the essential sources of food, nutrition, income and livelihood for people around the world. The efforts to promote aquaculture has been recently accelerated towards achieving the goal of food security. One of the recent efforts is to source sustainable plant proteins as an alternative to highly competitive and expensive fish meal and fish oil [^1^] for determining their nutritional suitability in fish diets [^2^]. The sustainable use of our aquatic resources can only be achieved through concerted efforts in using technology and human resources more effectively by enhancing our understanding about the changing nature of the aquaculture industry. The growth of world aquaculture has been stimulated by several factors, including population increases, dietary shifts and advances in aquaculture technology. Diet formulation involve the selection and mixing of feed ingredients to form products that delivers the much-needed nutrients to meet the production goals in fish farming. This practice can also reduce feeding time and feed waste while improving water quality and environmental safety of fish production.

Commercial feeds are expensive as most of the ingredients are imported and thus prices are increasing continually. So, finding alternatives to fish meal is needed to reduce cost and reliance on this valuable and finite source of protein. According to Hardy [3], one of the methods to develop less expensive and effective formulations is lowering fish meal levels in diets. Increased utilisation of plant protein meals has been embraced as a sustainable alternative to fish meal [^4, 5^]. Today, the most widely applied alternative plant protein to replace fish meal in aquaculture is soybean meal, mainly as extracted and protein concentrates [^6^], though with varied success. Several reports have suggested the partial substitution of soybean meal for fish meal in quantities for up to 60% or the application of its by-product [^1, 7, 7^]. The use of such a massive percentage in feed was possible due to their protein and amino acid contents, alongside digestibility and palatability properties [^9, 10^]. However, studies with culture fish have shown that soybean meal induces intestinal inflammation [^11^], which indicated that there was a need to research in other available plants sources (e.g. legumes) locally available with a reasonable amino acid profile that can replace fish meal. Plant protein sources like rapeseed meal, lupin seed meal, sunflower seed meal, peanut or cottonseed meal had also found their application, though with some limitations due to their anti-nutritional components. The rapeseed meal has high nutritional value, and thus great potential as an alternative protein source for fish nutrition [^12^]. The highest fishmeal replacement level of up to 100% have been reported in feeding trial with rainbow trout (*Oncorhynchus mykiss*), 30% substitution of fish meal protein for common carp (*Cyprinus carpio*) [^12^] and 25% substitution of fish meal protein for catfish [^13^] without affecting the growth performance of those fish. It is one of the world-leading edible oilseed crops. It has high protein content, which is distinguishable by a well-balanced amino acid composition and a high biological value [^14^]. The heat treatment of the plant protein reduces the anti-nutritional factors and increase the apparent protein digestibility [^15^].

Zebrafish (*Danio rerio*) formerly known as *Brachydanio rerio* is one of the approximately 45 *Danio* species that are known to exist all over the world [^16^]. They are tropical freshwater fish which belong to the minnows and carp family - Cyprinidae of the order Cypriniformes. It is native to Himalayan region and particularly indigenous to Ganges region of India, Pakistan, Bangladesh, Nepal and Burma and commonly inhabit the streams, canals, ditches, ponds, slow-moving or and stagnant water bodies including rice fields [^16-23^]. In the field study of zebrafish location, it was reportedly found to be common in shallow water bodies with turbidity level of up to about 30 cm, and most often in unshaded sites with open water among aquatic vegetation and silty substratum. Zebrafish (*Danio rerio*) has been used extensively in a number of scientific research disciplines (e.g. developmental biology and genetics; [^24-26^]; comparative biology [^19^]; toxicology [^27^], evolutionary theory [^28^], and aquaculture, immune response, nutrition and growth [^11, 17, 21, 29-31^]. However, the scope of their use as a model for nutrition and aquaculture research has been limited, partly due to lack of standardised or defined diets and husbandry conditions.

Many ingredients that are common components of aquatic diets contain compounds that alter the physiology and behaviour of an organism [^32^]. In the existing feeding procedures for *Danio rerio*, a few studies have evaluated the effects of diets on growth and survival of *Danio rerio* [^33^] indicating that the availability or unavailability of particular nutrient in the diets could affect normal physiological processes of fish. Also, Gonzales Jr and Law [34] showed that the use of commercial feeds and various feeding regimes affect the growth and gonadal development in zebrafish. Thus, the development and growth in fish are influenced by the changes in various dietary nutrients.

The standard feeding practice for zebrafish cited in numerous publications has been the administration of a diet which is primarily from a natural source –*Artemia nauphli* and a processed feed offered as tropical fish flakes when the fish grows older [^35-37^]. The common usage of flakes in zebrafish diets is problematic because such foods had been reported to contain potentially harmful; compounds such as genistein and a soy isoflavone which is shown to be an estrogen mimic in mammals and fish [^38, 39^]. The practices of feeding and feed management have numerous implications for the success of the normal zebrafish and aquaculture operation. Despite this fact, the optimal feeding frequencies for post-larval zebrafish have never been formally determined. Zebrafish is chosen for this study because of its many advantages over other teleosts. It is easy for handling and breeding experimentation, due to short generational intervals. They produce large numbers of eggs per brood (100-200 eggs/clutch) [^40^], which allows elaborative analysis with replicated number of specimens per data point and genetic manipulation [^41^].

The central hypothesis of this research was to test the efficiencies of plant-based diets as a positive replacement of commercial FM based diets on growth performance and body composition of zebrafish as a model organism. The study used repeated and non-invasive methods for morphometric measurement using a digital calliper and cameral imaging to determine the condition factor. The study compared SBM and RSM based diets with a commercial FM based diet for acceptability and effects on growth and body composition of zebrafish. It also evaluated the interactive effects of stocking density and the diets on growth and body measurement of *Danio rerio*.

## Material and Method

### The experimental fish, husbandry and design

The juvenile zebrafish (AB strain) were acquired from the zebrafish unit of Newcastle University. Fifty adult wild type zebrafish were used in a 1:1 ratio of a male to a female in each 1-L tank, where a total tank density of 2 fish per spawning tank was maintained to breed the fish. Each breeding tank consisted of a false-bottom tank containing a mesh screen bottom that was seated within a relatively larger tank to allow separation of released eggs and milt. The embryos were collected and rinsed with a 60-ppm saline solution containing 1ml methylene blue/L. This solution was used for cracking the eggs to facilitate hatching and removal of unfertilised eggs, and to prevent microbial infestation of the eggs. Fertilised embryos were placed in culture dishes at a density of 30-50 embryos per dish which were incubated at 28 °C. After five days post fertilisation (dpf), the larvae were placed in a 2-L tank and fed on commercial diets initially with a particle size of ZM-000 - 200 fry food for approximately six weeks and later on tropical fish flakes (ZM: Fish Food/www.ZM system.co.uk). At sixty-two dpf, fish were distributed into eighteen rectangular tanks (31.5cm x 13.5cm x 13.0cm) each with 3-L water capacity. The tanks were connected to a common water source, and an aquarium pump (Surpass air pump – Resun® Air-8000, China), which supplied oxygen continuously throughout the experiment to reduce the accumulation of nitrogenous waste in the tank water. Twelve fish were stocked in each of these eighteen tanks according to the 3 × 2 factorial arrangement. A light: dark photoperiod of 14:10 hours was provided with indirect fluorescent lighting. The water quality parameters, which include dissolved oxygen, pH, ammonia, water hardness, chlorine, nitrites (NO_2_) and nitrates (NO_3_), were determined weekly. Dissolved oxygen was measured by using commercial kit (CHEMetrics Inc. VA 22728, USA). The temperature was monitored daily using an aquarium thermometer (Pond planet, Cleveland, UK). The pH, ammonia, nitrate and nitrite, chlorine, and water hardness were monitored weekly using an aquarium Tetra 6 in 1 water test strip (Tetra GmbH, Germany). All fish tanks containing water were maintained at 26°C, pH 6.8 and 370 μS/cm conductivity in a recirculating system.

### The experimental diets and fish feeding

Three diets comprising one commercial diet (TetraMin Tropical Flakes® with 47% protein, 10% fat, 3% fibre; (Tetra Holding GmbH, Germany) and two plant protein-based diets containing either SBM or RSM were used in this study (Table 1). The feed ingredients were acquired from A-One Feed Supplements Ltd, Thirsk, North Hill, Dishforth Airfield YO7 3DH, North Yorkshire, England. Ground brown rice, cod liver oil, NaCl (salt) were purchased from Holland and Barrett, Nuneaton, Warwickshire, UK), Chromium III oxide (Cr_2_O_3_ was purchased from Sigma Aldrich, UK. The dried ingredients were ground to fine particles with a coffee grinder (DeLonghi KG49, UK) and thoroughly mixed by using a blender (KM 336, Kenwood, UK). The ground ingredients were sieved by using Laboratory test sieve with 250 μm aperture (281716, ENDECOTTS LTD, London, UK) to obtain uniform particle size for the juvenile fish diet.

**Table 1.** Ingredients and nutrient composition of the experimental diets.

| Ingredients | CFM diet | SBM diet (g/kg) | RSM diet (g/kg) |
| --- | --- | --- | --- |
| Fish Meal (64%) | - | 229 | 240 |
| Rape Seed Meal (37%) | - | - | 240 |
| Soybean Meal (50%) | - | 229 | - |
| Wheat gluten (60%) | - | 229 | 240 |
| Yellow Maize | - | 66 | 53 |
| Wheat Bran | - | 65 | 53 |
| Ground Brown Rice | - | 65 | 53 |
| Lysine (Lys) | - | 10 | 10 |
| Methionine | - | 12 | 12 |
| CaCO<sub>3</sub> (Ca<sup>2+</sup>) | - | 3 | 3 |
| Salt | - | 4 | 4 |
| Vitamin Premix | - | 15 | 15 |
| Mineral Premix | - | 10 | 10 |
| Cod-liver oil | - | 58 | 65 |
| (Cr<sub>2</sub>O<sub>3</sub>) | - | 5 | 5 |
| **Proximate composition of diets (g/kg)** |  |  |  |
| Dry matter | 928.94 ± 5.39 | 916.49 ± 8.87 | 925.83 ± 5.75 |
| Crude protein | 448.07 ± 2.45 | 474.81 ± 17.07 | 443.02 ± 3.02 |
| Crude fat | 125.88 ± 8.85 | 109.76 ± 3.49 | 143.44 ± 18.04 |
| Crude fibre | 31.54 ± 6.14 | 75.80 ± 6.58 | 110.62 ± 4.24 |
| Ash | 39.35 ± 0.45 | 68.48 ± 0.79 | 70.62 ± 0.67 |
| Nitrogen-free extract | 284.09 ± 4.68 | 187.65 ± 12.08 | 139.64 ± 7.42 |
| **Mineral composition (g/kg of mineral)** |  |  |  |
| Calcium | 49.96 ± 0.46 | 80.84 ± 0.55 | 85.29 ± 0.10 |
| Potassium | 13.97 ± 0.66 | 18.86 ± 0.62 | 10.50 ± 0.39 |
| Magnesium | 5.23 ± 0.04 | 5.97 ± 0.15 | 5.48 ± 0.06 |
| Sodium | 16.06 ± 0.21 | 1277 ± 0.20 | 14.05 ± 0.14 |
| Phosphorus | 2.01 ± 0.04 | 2.00 ± 0.02 | 1.94 ± 0.02 |
| Zinc | 0.32 ± 0.00 | 2.21 ± 0.02 | 2.56 ± 0.04 |
| Iron | 1.12 ± 0.00 | 4.01 ± 0.03 | 3.73 ± 0.00 |
| **Essential amino acids composition (g/kg of protein)** |  |  |  |
| Histidine | 7.40 ± 1.30 | 3.78 ± 0.09 | 3.85 ± 0.20 |
| Leucine | 50.73 ± 0.90 | 66.55 ± 2.76 | 68.12 ± 3.69 |
| Isoleucine | 24.06 ± 1.30 | 21.62 ± 0.47 | 17.91 ± 1.95 |
| Lysine | 62.44 ± 14.68 | 33.18 ± 1.03 | 30.35 ± 1.42 |
| Methionine | 5.48 ± 0.53 | 11.63 ± 1.14 | 12.46 ± 0.64 |
| Phenylalanine | 50.49 ± 2.63 | 40.31 ± 6.24 | 39.32 ± 3.50 |
| Threonine | 2.54 ± 0.41 | 2.60 ± 0.24 | 2.54 ± 0.34 |
| Valine | 13.54 ± 1.13 | 11.94 ± 0.38 | 10.03 ± 1.30 |

Representative samples of the diets were analysed for their proximate composition, minerals contents and fatty acid profile. About 100g of the ingredient mixture was made into a slurry by mixing with about 250 ml warm water. This semi-liquid mixture was thinly spread inside aluminium foil trays before freezing at −20 °C followed by freeze-drying for five days to a constant weight. Feeding of the experimental fish was accomplished manually. The fish were fed a quantity of feed worth 5 % of their body weight. Feeding of the fish was carried out three times daily (6 – 8 am, 12 – 2 pm and 6 – 8 pm) by following the published guidelines [^42, 43^]. The amount of feed administered was adjusted based on the weight of the fish in each weak. However, overfeeding was avoided to prevent wastage, water contamination and possible microbial growth and infection.

### Morphometric measurements of fish

Representative samples of fish in each group were assessed weekly for their length and weight. This assessment was accomplished by weighing all the fish in a tank as a unit to the nearest 0.01mg. One fish was randomly selected from each tank and euthanised with tricaine methanesulphonate MS-222 (Sigma) in 100 ml aqueous solution at 200 mg/L as used by Matthews, Trevarrow [35] for morphometric measurement. Each euthanised fish was then placed in a petri dish with a clear background and photographed with a digital camera (Samsung ES90). Image J and MATLAB computer software were used to obtain the measurement of both length and width from the captured images with the camera. A digital calliper (Silverline®, 380244, London UK) was also used to measure the length and width of the fish. All length measurements were recorded to the nearest 0.01mm. The tail and mouth ends were acquired as optimal points to describe the total length of the fish as the fish share the same genetic characteristics of their morphology. At the end of the experiment (118 dpf), each fish was euthanised and then measured for its weight and length. The measurements of both length and weight were used to determine the condition factor (the well-being of the fish) according to Fulton condition factor [^44^].

### Growth performance

Various measurements and growth performance indices were evaluated. The weight gain was calculated as the final weight (g) – the initial weight of the fish per tank. MWG/Day = mean weight gain (g) / number of days. The specific growth rate (SGR) = (ln w_2_ - lnw_1_) / (t_2_ - t_1_) x 100. Where: W_1_ = weight of fish at the beginning of the experiment (t_1_ days), W_2_ = weight of fish at the end of the experiment (t_2_ days), ln = natural logarithm and t_2_ – t_1_ = experimenter periods in days. Total feed intake was calculated as the cumulative food consumed by each fish group per tank. The FCR was calculated according to Houlihan, Boujard [^45^] from the relationship between food intake and weight gain by fish. (i.e.) FCR = food intake (g) / weight gain (g). The protein efficiency ratio (PER) = net weight gain (g) / protein intake in term of the nitrogen content of the feed (g). Daily protein intake (g) was computed by the relationship between the protein intake and the number of days of the experiment. DPI = protein intake/number of days. The gross energy was determined by using an adiabatic bomb calorimetric (Gallenkamp auto bomb, CBA-305-010M, UK) with a benzoic acid standard. The result was cross-referenced with energy determination using the conversion factor [^46^]. The mathematical formula that determined condition factor according to Fulton condition factor (Fulton, 1902), was as follow: K_FL_ = W/L^3^ x 100, where W= fish weight (mg), and L= fish total length (mm) while K is the coefficient of condition.

### Proximate composition

The proximate composition of the freeze dried samples of fish was performed by the application of AOAC [^47^] methods. The freeze-dried fish samples were grounded with a coffee grinder (DeLonghi Living innovation, UK). Moisture content and the dry matter were determined after freeze-drying when a constant weight was obtained. The nitrogen (N) and carbon (C) contents of the fish tissue were determined by using the cube method with Elementar machine (Elementar Vario Macro cube, Germany). The crude protein was evaluated by multiplying the N with the conversion factor of 6.25 [^46^]. The fat content was extracted by using dichloromethane and methanol (2:1 *v/v*) (98.9%) (Sigma Aldrich, UK) in a modified method of Folch, Lees [48]. The ash content was determined after combustion of the sample in a muffle furnace at 550 °C for five hours. All the analysis was conducted in triplicates.

### Statistical Analysis

The data collected were subjected to normality test to ascertain if the data was normally distributed or not using IBM SPSS Statistics for Windows, Version 22.0. Armonk, NY: IBM Corp. The data that were not significantly different from normal distribution were subjected to analysis of variance (ANOVA) and two samples Student T-test. The effects of the diets, stocking density and the interaction of the diet with the stocking density were determined by GLM (general linear model). The effects of stocking density and diet on weekly growth were studied. The Mauchly’s test of sphericity and the Levene’s test of equality of variation were used to analyse the differences in the growth of the fish. Tukey’s post-hoc test was used to compare mean to observe significance at p < 0.05.

## Results

The results of the percentage survival of the fish at the end of the experiment are presented in Figure 1. The percentage of survival ranged between 95.8 % in the fish fed SBM diet to 97.2 % in the fish fed CFM and RSM diets, respectively. The high survival rate of zebrafish fed the plant protein-based diet compared to those fed commercial FM-based diet revealed that the feeding zebrafish with a high percentage of plant protein-based diet was possible without causing any adverse effects on their survival.

**Figure 1.**
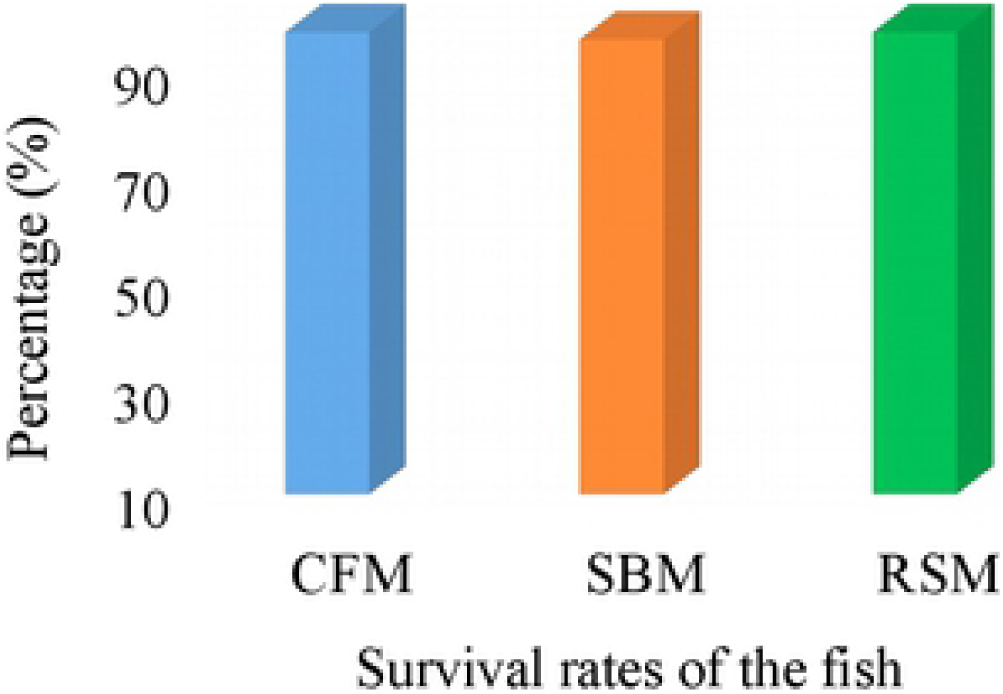
The survival rates of the fish fed the commercial fishmeal (CFM) or soybean rneal (SBM) or rapeseed meal (RSM) diets, respectively.

### Morphometric measurements of zebrafish

The values of the mean and standard errors for the maximum length and weight attained by the fish at the end of the feeding trial for each of the diets and the stocking density are presented in Table 2, and 3 respectively. Fish fed the commercial diet had significantly higher (P<0.05) length and weight (32.54 ± 0.42mm and 449.75 ± 22.83mg) than the fish fed either SBM or RSM based diets. However, there was no significant difference (p>0.05) in both length, and weight between the fish fed RSM (30.01 ± 0.43mm, 329.25 ± 19.82mg) and SBM diets (30.23 ± 0.36mm, 329.69 ± 60.06mg). There was no significant difference between the stocking density for the fish length and weight, though the mean weight and length of fish were numerically higher for the fish stocked at lower stocking density than those stocked at higher density. The results of the length-weight relationship of the fish presented in Figure 2 show that the fish fed the respective diets exhibited a positive correlation between the length and the weight during the experiment. It shows the coefficient of determination R-sq and the regression equation for the respective dietary factor where CFM (n=55), SBM (n=56), and RSM (n=5).

**Figure 2.**
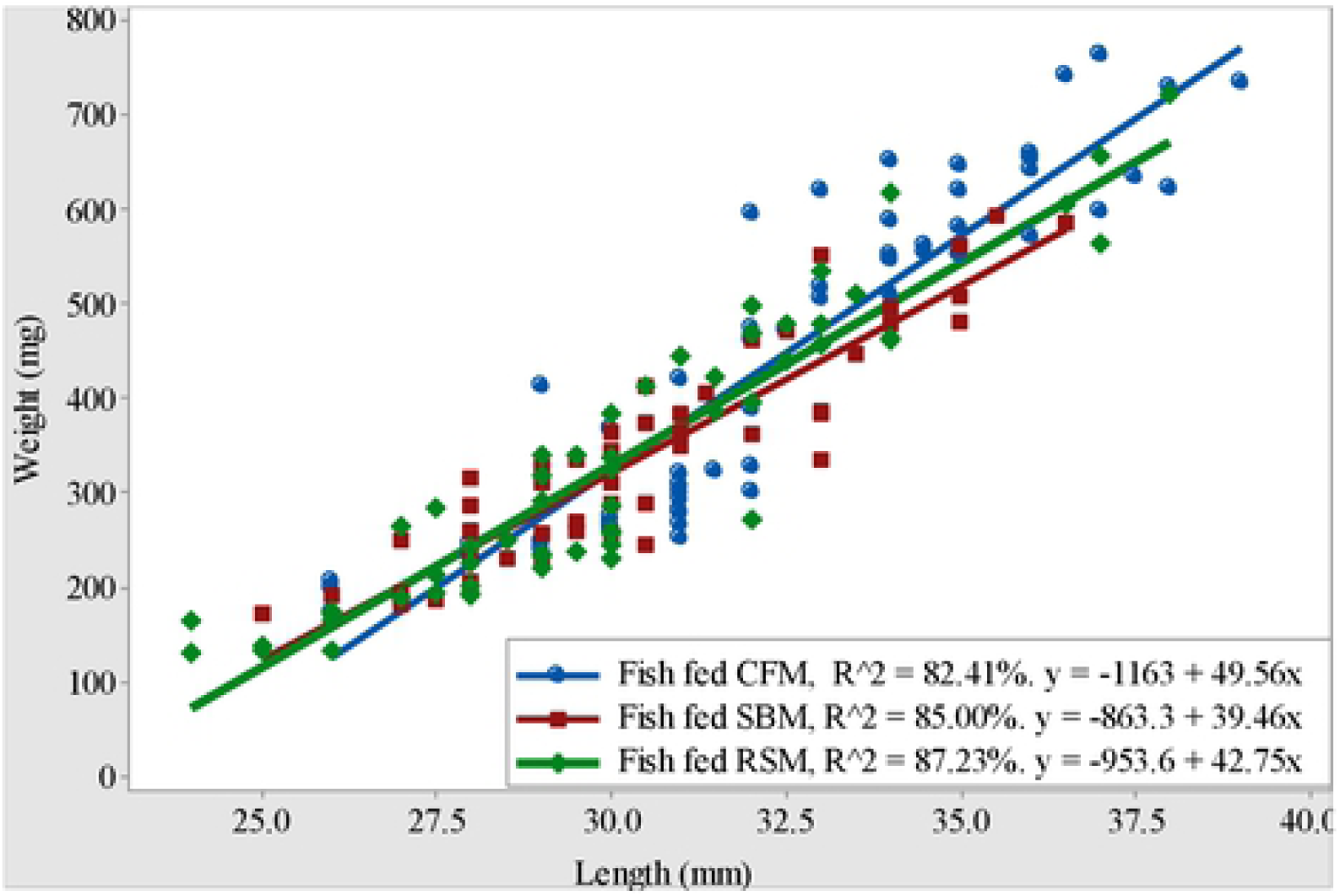
Scattered diagrams with fitted lines of the length-weight relationships of the fish fed the various diets.

**Table 2.** Mean length and weight of the fish for the main effects of diets and the stocking density (total fish)

| Variables | CFM<br>(n = 56) | SBM<br>(n = 55) | RSM<br>(n = 56) | Total fish NS<br>(n = 104) | Total fish RS<br>(n = 63) |
| --- | --- | --- | --- | --- | --- |
| Wt. (mg) | $449.75 \pm 22.8^a$ | $329.69 \pm 15.5^b$ | $329.25 \pm 19.8^b$ | $359.51 \pm 15.7$ | $386.79 \pm 18.9$ |
| *P - values* | 0.000 |  |  | 0.26 |  |
| L (mm) | $32.54 \pm 0.42^a$ | $30.23 \pm 0.36^b$ | $30.01 \pm 0.43^b$ | $30.73 \pm 0.33$ | $31.26 \pm 0.36$ |
| *P - values* | 0.000 |  |  | 0.283 |  |

The mean values with the same superscript along the same row for diets and the stocking density are not significantly different ( $p > 0.05$ ). NS = high fish stocking density and RS = low fish stocking density.

**Table 3.** Mean length and weight of the zebrafish for the various diets and stocking density interaction.

| Variables | CFM |  | SBM |  | RSM |  |
| --- | --- | --- | --- | --- | --- | --- |
|  | NS<br>(n = 35) | RS<br>(n = 21) | NS<br>(n = 34) | RS<br>(n = 21) | NS<br>(n = 35) | RS<br>(n = 21) |
| Wt. (mg) | 439.9 ± 29.8 | 466.2 ± 35.8 | 324.7 ± 21.7 | 337.9 ± 21.0 | 313.0 ± 24.5 | 356.3 ± 33.5 |
| *P-Values* | 0.574 |  | 0.663 |  | 0.303 |  |
| L (mm) | 32.41 ± 0.54 | 32.74 ± 0.67 | 30.14 ± 0.51 | 30.38 ± 0.48 | 29.61 ± 0.58 | 30.67 ± 0.62 |
| *P-Values* | 0.708 |  | 0.736 |  | 0.222 |  |

The mean values with the same superscript along the same row for diets and the stocking density are not significantly different ( $p > 0.05$ ). NS = high stocking density and RS = low stocking density. CFM, SBM, and RSM are commercial fishmeal, soybean meal, and rapeseed meal-based diets, respectively

The regression coefficient (R^2^) was highest in fish fed RSM diet (87.23 %) compared to that of the SBM diet (85. 00 %) and CFM (82.41 %) respectively. There were significant correlations between the body weight and the length for the three dietary treatments ranging from (r = 0.91) in fish fed CFM diet to (r = 0.93) in fish fed RSM diet.

The mean condition factors were 1.2 ± 0.2, 1.15 ± 0.2 and 1.14 ± 0.2 for the fish fed CFM, SBM, and RSM based diet, respectively. The results of the Fulton condition factor show that all the fish fed the three diets exhibited positive allometric condition indicated by the growth coefficient (b>3). The value is an indication that the fish grows faster in their weight than in their length. The state of the well-being of the fish fed the diets were good as indicated by the calculated condition factor (k>1) (Figure 3). The n = 56, 55, and 56 for the fish fed CFM, SBM, and RSM diets respectively. There was no significant difference in the condition factor (K) of the fish fed the various diets.

**Figure 3.**
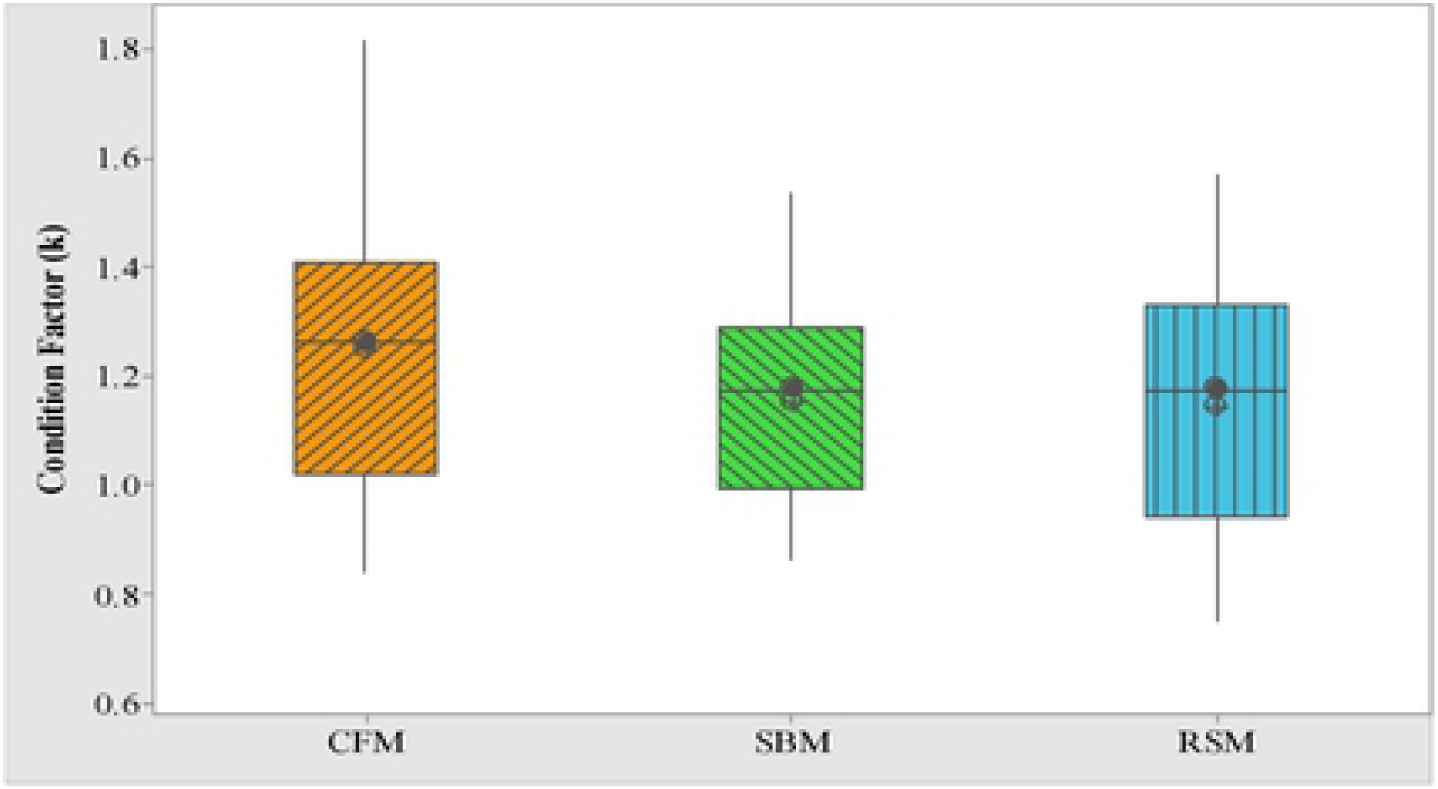
Box plot showing the mean and median values of the condition factor of the fish fed the three different diets.

### Growth performances of zebrafish fed the three diets

The results of the weekly growth pattern based on the three diets and the two stocking densities are presented in Figure 4. The fish fed the CFM diet recorded the highest growth. All the fish exhibited a similar pattern of growth irrespective of the diet fed. At the initial stage, the weight dropped in the fish fed SBM and RSM diets (Fig. 4 B & C). After acclimation to the new diets and the environment, the growth patterns of the fish changed. A similar trend was observed in tanks with low and high fish stocking density among the fish fed the three diets.

**Figure 4.**
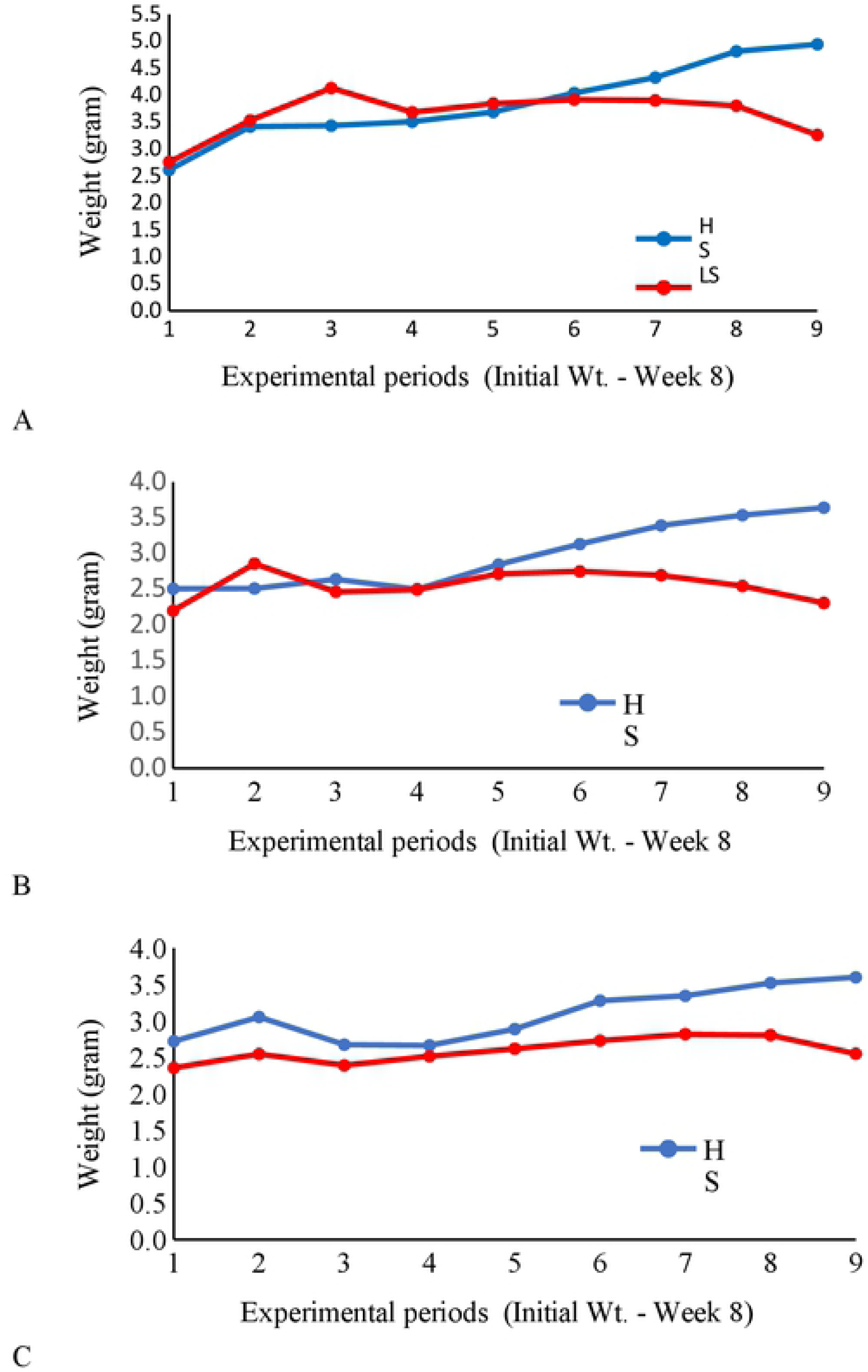
The weight measurement of the fish fed various diets weekly at two stocking densities. A is the fish fed commercial fish meal-based diet, B is the fish fed soybean meal-based diet, and C is the fish fed rapeseed meal-based diet. HS represents high stocking density and LS stand for low stocking density. 1, is the initial weight of fish while 2-9 are the weights of fish for eight weeks.

The multivariate test results show that there were significant differences (p<0.05) in the weight of the fish based on the stocking density. However, there was no significant difference observed among the fish fed the diets. The Mauchly’s test revealed no significant differences in the weight of the fish relative to the interaction between the diets and the stocking density. The Mauchly’s test of sphericity equally revealed significant differences (p<0.05) in the weight of the fish between the first week and the second week of the feeding trial. There were no significant differences; however, in the weight of the fish between the second, and the third week until the fourth week through to the seventh week where significant increments (p<0.05) were observed. Significant differences were manifested in the weight of the fish between the two stocking densities from the fifth weeks to the eight weeks. The difference was due to the reduction in the numbers of the fish due to the fish sampling relative to the respective culture medium. Levene’s test of equality of error of variances across the groups from one week to the other showed that there were no significant differences (p>0.05) in the error variances of the weight of the fish. On the overall performance of the fish, the results show a significant interaction of the diet and the stocking density on the weight gain, specific growth rates, food conversion ratio, and the protein efficiency ratios of the fish during the period (Figure 5).

**Figure 5.**
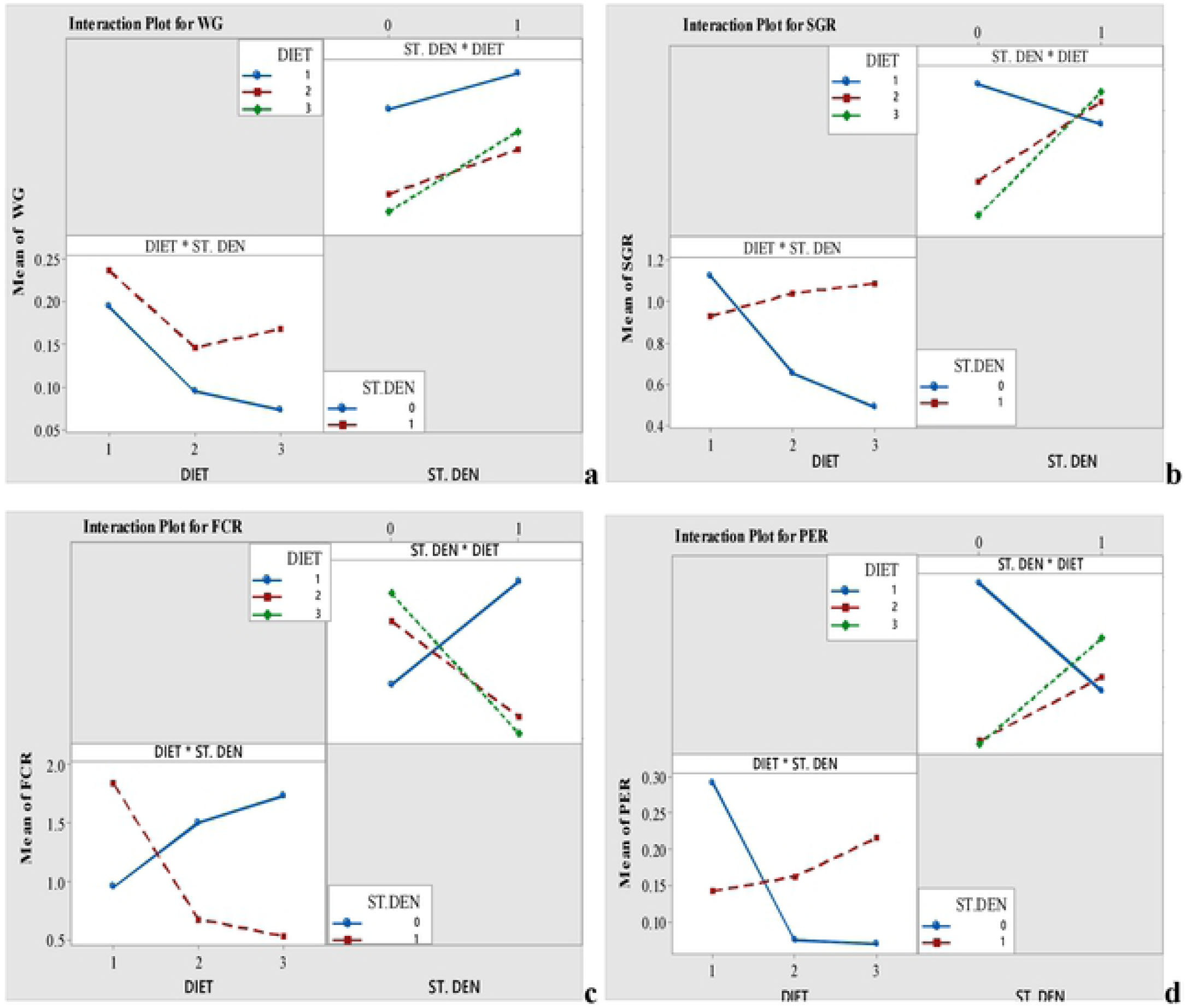
The interaction of the diet and stocking density on some specific growth parameters of zebrafish fed commercial fish meal and plant protein-based diets.

### Nutrients utilisation and growth performance

Table 4 and 5 show the results of the growth performance of the fish fed the three diets, and the respective stocking densities. There were significant differences (p<0.05) in the weight gain (WG), protein intake (PI) and feed conversion ratio (FCR) among the fish fed CFM, SBM and RSM diets. The values were significantly higher in the fish fed the CFM than the other two diets. However, there were no significant differences observed (p>0.05) in the SGR, FI and the PER in the fish fed the three diets. The fish fed CFM diet converted more food to flesh and gained more weight than those fed other diets. Likewise, fish under the low stocking density gained more weight significantly (p<0.05) than the fish in tanks with high stocking density. The PI was significantly higher in the fish stocked at high density than those at low density. There were no significant differences (p>0.05) in the SGR, FCR, FI, as well as the PER between the two stocking densities among the fish fed the three diets. Fish in tanks with low stocking density converted more food to flesh and gained more weight than those stocked at high density. There were no significant differences between the two stocking densities in the fish fed CFM diet with respects to all the growth indices except the weight. However, the results were different in the fish fed either SBM or RSM diets. The WG was similar in both stocking densities for the fish fed both SBM and RSM based diets. The PER and SGR were higher for fish with low stocking density among those fed RSM and SBM diets. However, FI, FCR, and PI were not significantly different (p>0.05) in fish relative to both high and low stock densities.

**Table 4.** Mean growth performance indices of the experimental fish

| Variables | CFM<br>(*n*=6) | SBM<br>(*n*=6) | RSM<br>(*n*=6) | Total fish |  |
| --- | --- | --- | --- | --- | --- |
|  |  |  |  | HS (*n*=9) | LS (*n*=9) |
| Ini. Wt (g) | 2.11 ± 0.25 | 1.90 ± 0.29 | 2.05 ± 0.31 | 2.61 ± 0.09<sup>a</sup> | 1.42 ± 0.07<sup>b</sup> |
| Fin. Wt (g) | 4.10 ± 0.40 | 2.98 ± 0.32 | 3.07 ± 0.27 | 4.06 ± 0.25<sup>a</sup> | 2.71 ± 0.16<sup>b</sup> |
| WG (g) | 1.99 ± 0.16<sup>a</sup> | 1.08 ± 0.06<sup>b</sup> | 1.03 ± 0.11<sup>b</sup> | 1.45 ± 0.23 | 1.28 ± 0.10 |
| FI (g/d) | 0.17 ± 0.01<sup>a</sup> | 0.12 ± 0.01<sup>b</sup> | 0.11 ± 0.01<sup>b</sup> | 0.15 ± 0.01 | 0.12 ± 0.01 |
| FCR | 0.09 ± 0.01 | 0.12 ± 0.01 | 0.11 ± 0.01 | 0.12 ± 0.01 | 0.09 ± 0.01 |
| SGR (%/d) | 1.18 ± 0.06<sup>a</sup> | 0.85 ± 0.10<sup>ab</sup> | 0.79 ± 0.14<sup>b</sup> | 0.76 ± 0.10<sup>b</sup> | 1.13 ± 0.05<sup>a</sup> |
| PI (g) | 1.45 ± 0.11<sup>a</sup> | 1.12 ± 0.08<sup>ab</sup> | 0.92 ± 0.09<sup>b</sup> | 1.29 ± 0.09 | 1.03 ± 0.10 |
| PER | 1.38 ± 0.07 | 0.99 ± 0.08 | 1.17 ± 0.17 | 1.07 ± 0.11 | 1.29 ± 0.09 |

Values are means ± standard error. The means with the same superscript along the same row are not significantly different at ( $p>0.05$ ). HS = High stocking density, LS = Low stocking density, FCR = Food conversion ratio, SGR = specific growth rate, NA = Not applicable 'n' = the number of replicates. CFM, SBM, and RSM are the fish fed the various diets. There are 56, 55, and 56 fish samples for each of the six replicates under each dietary treatment, respectively.

**Table 5.** Mean growth performance indices among the fish fed the various diets based on the stocking density, and (Diet x stocking density interaction)

| Variables | CFM |  | SBM |  | RSM |  |
| --- | --- | --- | --- | --- | --- | --- |
| | HS ($n = 3$) | LS ($n = 3$) | HS ($n = 3$) | LS ($n = 3$) | HS ($n = 3$) | LS ($n = 3$) |
| I. Wt (g) | $2.61 \pm 0.21^a$ | $1.61 \pm 0.09^b$ | $2.51 \pm 0.17^a$ | $1.28 \pm 0.13^b$ | $2.72 \pm 0.14^a$ | $1.37 \pm 0.11^b$ |
| F.Wt (g) | $4.94 \pm 0.26^a$ | $3.26 \pm 0.06^b$ | $3.64 \pm 0.21^a$ | $2.31 \pm 0.12^b$ | $3.60 \pm 0.24^a$ | $2.55 \pm 0.16^b$ |
| Wt.G (g) | $2.33 \pm 0.05^a$ | $1.65 \pm 0.10^b$ | $1.13 \pm 0.14$ | $1.03 \pm 0.03$ | $0.88 \pm 0.01$ | $1.17 \pm 0.01$ |
| SGR (%) | $1.13 \pm 0.05$ | $1.25 \pm 0.10$ | $0.66 \pm 0.08^b$ | $1.04 \pm 0.09^a$ | $0.49 \pm 0.08^b$ | $1.09 \pm 0.05^a$ |
| F.I (g) | $0.19 \pm 0.01$ | $0.16 \pm 0.02$ | $0.15 \pm 0.00$ | $0.11 \pm 0.01$ | $0.12 \pm 0.01$ | $0.09 \pm 0.01$ |
| FCR | $0.90 \pm 0.06$ | $0.66 \pm 0.07$ | $1.36 \pm 0.32$ | $0.74 \pm 0.09$ | $0.15 \pm 0.04$ | $0.08 \pm 0.01$ |
| PI (g) | $1.59 \pm 0.09$ | $1.31 \pm 0.17$ | $1.26 \pm 0.04$ | $0.99 \pm 0.13$ | $1.04 \pm 0.10$ | $0.80 \pm 0.12$ |
| PER | $1.48 \pm 0.06$ | $1.29 \pm 0.12$ | $0.90 \pm 0.09$ | $1.07 \pm 0.13$ | $0.84 \pm 0.12^b$ | $1.51 \pm 0.15^a$ |

Values are mean ± standard error. The mean values with the same superscript in the same row under the same diet are not significantly different ( $p>0.05$ ). HS = high stock density, LS = low stock density, n = the number of replicates. I.Wt and F.Wt = the initial and final weight, SGR = specific growth rates, F.I = the food intake, FCR = the food conversion ratio, PI = protein intake, and PER = protein efficiency ratio, while CFM, SBM and RSM are the fish fed the various diets respectively.

### The chemical composition of the fish fed the three diets (g/kg DM)

The proximate composition of the fish fed the CFM, SBM and RSM based diets are presented in Figure 6. There were no significant differences (p>0.05) in the dry matter (259.22 ± 18.02, 230.53 ± 16.59 and 265.30 ± 13.61), crude protein (137.76 ± 9.90, 131.9 ± 9.78 and 144.96 ± 7.63), ash contents (10.92 ± 0.32, 1.20 ± 0.28 and 1.61 ± 0.22) and the nitrogen-free extract contents (3.05 ± 0.31, 3.04 ± 0.28 and 2.95 ± 0.42) for the fish fed CFM, SBM and RSM respectively. However, the ether extract of the fish fed SBM was significantly lower (5.70 ± 0.50) than that of the CFM and RSM (8.46 ± 0.61 and 7.35 ± 0.60 respectively).

**Figure 6.**
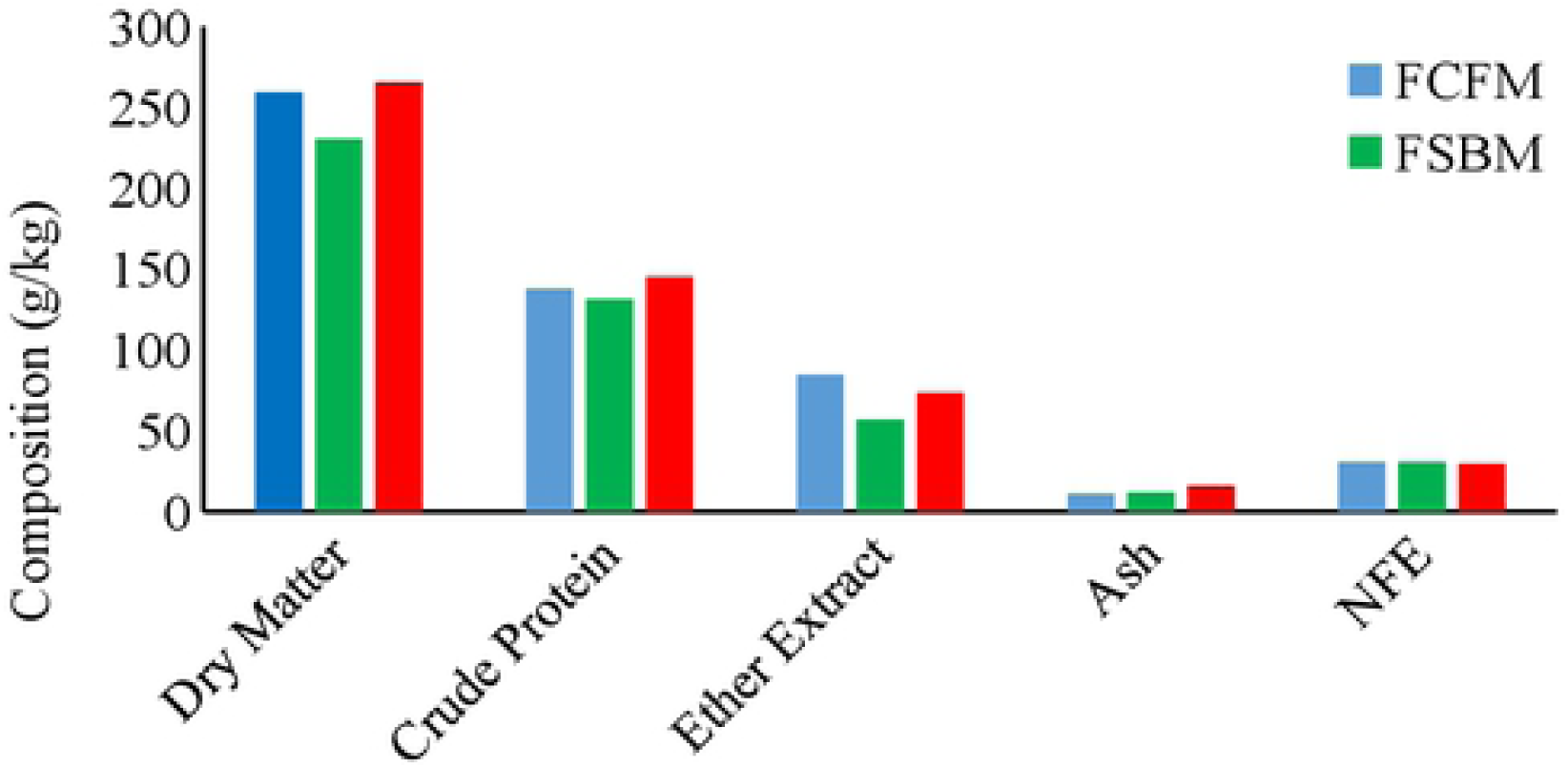
The chemical compositions (g/Kg DM) of the fish fed CFM, SBM, and RSM based diets. Data for crude protein, ether extract, ash, and NFE are based on the DM. NFE = Nitrogen free extract. Bars with the same letter are not significantly different (p>0.05). The fat content was lowest in the fish fed SBM.

## Discussion

The fundamental approach to establishing a standard diet for any fish species as a model organism begins with several tests of diet acceptability, especially when the formulation is comparison-based with an expensive fish meal and fish by-products. Fish, accept a feed based on its availability, chemical and physical characteristics, the nutritional value, the environment, stressors and social interactions, and method of food delivery [^17, 45, 45^]. Various researchers have shown the possibility and recommendation of cost-effective plant protein sources for the formulation of feeds for tropical fish culture with varying rates of survival [^50-54^]. The survival rates of zebrafish in this study confirmed it as a euryphagous species with the ability to consume wide varieties of available food substances and convert those into body flesh [^40, 55^]. The results are similar to the finding of Silva-Carrillo, Hernández [56], who reported a survival rate above 95% for juvenile spotted rose snapper *Lutjanus guttatus* (Steindachner, 1869) fed SBM. Al-Thobaiti, Al-Ghanim [^50^] reported about 50 −100 % survival for Tilapia (*Oreochromis niloticus*) fed plant protein-based diets. A survival rate of 87 % was reported for zebrafish fed compound feed from the first feeding onward [^57^]. The survival rates of zebrafish in this study were higher than the mean survival ranging from 55 – 86 % reported at 27 dpf [^58^], and 76 % for zebrafish fed TetraMin Tropical flakes [^32^]. The lack of adverse effects on the survival of zebrafish when fed both SBM and RSM based diets agree with the findings of Siccardi III, Garris [^32^] and Smith Jr, Barry [^59^] who observed no significant differences in the overall health condition and survival of *Danio rerio* fed different diets as determined by the analysis of their condition factor, which is classically considered a health and well-being indicator in fish.

Accomplishing measurement of fish in fisheries study and biological research is a challenging task but it is a beneficial instrument in determining the size of fish as it grows. Such assessment provides a guiding decision on grading, harvesting, marketing [^60^], developing management and conservation strategies [^61^], alongside policymaking objectives in fisheries [^62^]. The application of camera imaging in this feeding experiment is encouraging as this offers a fast and less invasive techniques for its use on a bigger scale for the future of aquaculture practices. In this study, the allometric growth (b <3>) (derived from regression coefficient) exhibited by the fish were similar to the findings of Nehemia, Maganira [63], and Ayo-Olalusi [64] on Tilapia species and African mud catfish reared in freshwater but those had lower condition factor. Balai, Sharma [65] reported similar negative allometric (2.60), isometric (3.02) and positive allometric growth (3.16) for *C. catla, L. rohita*, and *C. mrigala*, respectively. A similar finding was reported by Abobi [66] for some freshwater fish species from the White Volta, Ghana. The study further revealed good water quality and adequate feeding of zebrafish can promote better condition of the fish in a culture medium. The regression coefficient (b) was increased as the fish grew in this study which indicated that zebrafish like other tropical fish transit from negative allometric growth to positive allometric growth which can be attributed to age, size, and species [^66^].

Many environmental factors such as temperature, stocking density, and water quality as well as the genetic factors affect the growth and maturity of fish [^67-70^]. Besides, fish growth is influenced by body weight and feed consumption alongside the precocity of distinct sexes of fish observable in the growth of different tissues and physiological state of the fish [^68, 71^].

Growth is a quantitative trait which may be measured at different stages of an animal’s life cycle [^30^], with many variations occurring in the growth rates during laboratory experimentation. Several studies have been conducted to assess growth of zebrafish relative to nutrition and maturity [^69^], influence of protein source on body composition [^59^], growth and survival [^32^], influence of diets on eggs and larvae quality [^72^], nutrition and health [^73^], and nutritional requirement [^74, 75^].

The increased weight of zebrafish fed various experimental diet was an indication of diet acceptance by the fish and its ability to convert dietary protein into flesh. The differences observed in the growth indices could be attributed to the variations in the specific source of nutrients between the diets. These could have affected the physiological or molecular mechanisms that influence the weight of the fish fed the undefined FM-based diet [^32^] and longer habituation of the fish to the commercial diet than the other groups fed either SBM or RSM based diets. The less weight gain obtained in the fish fed RSM and SBM than the CFM based diets could also be attributed to their naturally occurring anti-nutritional factors and high fibre contents [^76-78^], that could have affected the nutrient utilisation and fish growth [^14, 79, 79^]. Estimated values of 5-16 g tannin/kg DM has been reported in the ingredient used in animal feeds. The value is highest in oilseeds, followed by legumes [^81, 82^]. This factor might have played an essential role in the reduced growth rate observed in the fish fed both SBM and RSM diets. The high feed intake might have also contributed to the weight differences between the treatments as fish were fed based on their body weight. The growth and FCR of fish are the leading indices for the determination of the acceptance and consumption of artificial diets. The FCR is mostly influenced by the visceromatic index (VSI), homogeneity, genetic factor, feeding, and well-being of a fish [^49, 83, 83^]. The FCR of fish and other aquatic smaller animals are affected by many factors among which are the water quality, rate and manner of feed presentation to the fish, and health status of the fish [^85^]. None of these factors was significant enough to affect the FCR in this present study. It measures averagely 1.4-1.8 for omnivorous species. The FCR recorded in this study were low compared to the reports of 2.8 to 1.2 for Salmon [^83^], lower values ranging from 0.9 to 1.3 were reported for Trout, Salmon, and Tilapia [^86^]. The low values compared to the recommended can be attributed to the size of the fish, high protein digestibility, and highly improved culture management [^83^]. The specific growth rates are one of the most frequently used functions in fisheries and aquaculture for biological interpretation of the growth per day. The SGR in this study was similar to the results obtained by Chambel, Severiano [87] who reported variable impacts of stocking density and diets on the SGR, condition factor and fish survival.

The lack of significant variation in the nutrient utilisation between the two stocking densities could be attributed to the schooling behaviour of zebrafish [^88, 89^]. This study showed that SBM and RSM fed fish at lower density gained more weight, had higher SGR, better FCR, and PER than those at high density. The study agrees with the findings of Vargesson [90] on zebrafish, Oppedal, Vågseth [91] on *Salmon salar*, Alhassan, Abarike [92] and Costa, Roubach [93] on *Oreochromis niloticus*, who reported that stocking densities influenced the growth and performances of fish. The study revealed that diet and stocking density played a vital role in the specific growth performance of zebrafish. The density-dependent growth regulation and biomass do occur in the late juvenile or adult stage of fish in aquaculture. The few weeks of culture or experimentation of this study may not be sufficient to conclude the effect of stocking density on zebrafish culture as remarked by Cowan et al., (2000), cited by [^94^], because the effects of stocking density manifested more as the fish grow bigger in aquaculture. The overall optimum growth performance of the fish fed the plant-based protein-based diet can be attributed to the high protein quality of SBM and RSM diets which compared favourably with the commercial FM based diets.

Although recent attention has been given to zebrafish for its potential as a model for genetics and different diseases in human beings, few studies have reported its body composition outcome relative to different diets. Various studies have demonstrated and quantified the body composition of fish adopted different approaches based on the environment, fish species and technologies. For example, Smith Jr, Barry [59] applied quantitative magnetic resonance (QMR) to establish body composition based on the lean and fat mass of zebrafish. González, Flick [95] studied yellow perch; Bolasina and Fenucci [96] used Brazilian codling (*Urophycis brasiliensis* Kaup, 1858); Chukwu and Shaba [97]; Siddique, Mojumder [98] studied marine fishes; and Suseno, Syari [99] used pelagic fish (*Amblygaster sirm* and *Sardinella gibbosa*); all these studies applied AOAC methods. Also, Breck [100] reviewed the body composition of fish from literature and generated a linear functional relationship between fish body lipid and body water. This present study applied the AOAC [47] methods to investigate the effect of SBM and RSM based diets on the body composition of zebrafish. The results in this study were in the range reported for various tropical fish species [^57, 101-107^]. However, the CP in this study was lower than that reported for *Clarias gariepinus* (18.34 - 20.32 %) and Tilapia species (18.65 – 18.74%) [^108, 109^] respectively. It was evident from the study that the diet affected the protein composition of the zebrafish.

The fish muscle constitutes 50-60 % muscle [^110^] of which the protein constituent is about 16 - 21 % and lipids of about (0.5 – 2.3 %). The ether extract recorded in this study placed zebrafish as high-fat fish (despite being a small fish) compared to the findings of Mazumder, Rahman [105] for some small fishes of Bangladesh. According to Coppes Petricorena [110], lipid and water are connected components in fish: hence, as lipid increases, the moisture content decreases as equally observed in this study. The varied composition recorded for different species relative to this study may be attributed to the environment and feeding factors [^104, 111^]. The results of the ash content of zebrafish in this study were similar to the report of Begum, Akter [112] and Bernardes, Navarro [103]. It is also in the range reported by Bogard, Thilsted [104], but lower than the findings of Njinkoue, Gouado [113] who reported 7.17 and 7.28 % ash contents for *Pseudotolithus typus* and *Pseudotolithus elongates* respectively. The diets had no significant effect on the protein and the carbohydrate (NFE) levels of the fish but significantly impacted the fat (FA), and ash (mineral) composition of the fish. The results could be attributed to the low-fat content of the SBM based diet in the feed compared to other ingredients. The fish body is constituted by four significant components often determined in proximate analysis viz: water (moisture), lipids (FA), ash (minerals), and protein (AA). These components constitute about 96 – 98 % of the total tissue of fish. Growth of fish is much related to these four-component alongside the carbohydrates, vitamins, nucleotides and other non-protein nitrogenous compounds which were present in variable and small quantities but played vital roles in the body maintenance, growth, and development of the organisms.

### Conclusion and recommendation

In general, the results of this study agreed with previous findings that zebrafish could digest plant protein-based diets for their utilisation and growth without adverse effects of the diets on the fish growth. However, the proximate composition of the whole fish body was, in most cases, affected by the dietary treatment (especially on the fat contents). The study showed that zebrafish could be used for morphometric studies as a model organism for aquaculture species. There was no information on the detailed body composition of zebrafish fed plant protein-based diets compared with the commercial diets used in most zebrafish laboratories; therefore, the results of this study will contribute to this invaluable database for zebrafish nutrition. An aquaculture system using zebrafish for nutrition research needs to develop a method to carry out and elucidate the interactive effect of different nutrient sources on feed digestibility and its effects on the growth and nutrient utilisation of fish. There is a need to determine the level of inclusion of plant protein-based diets for the formulation of a standard diet for zebrafish. Studies have shown that the protein content of up to 30-40 % or less in place of FM in the diets of some species could be used without adverse effect on their growth performance. However, the maximum levels of inclusion of plant proteins together with the mechanism involved in the dietary changes including digestibility and, anti-nutritional factors in a molecular base need further research to be able to adopt zebrafish as a model organism for aquaculture. Nutrition experiment with small size and slow growing zebrafish need a more extended dietary trial for the manifestation of visible changes that can be accounted for by the test factors in clarifying their effects on growth performance, feed efficiency and the body compositions. This study showed that both SBM and RSM diets could be used to replace high-cost CFM diet in the culture of zebrafish and other similar fish species in aquaculture.

## Acknowledgement

This research was supported by the PhD studentship funded by the Tertiary Education Trust fund and Ekiti Sstate University, Nigeria.

## Disclosure Statement

No conflict of interest exists.

